# Time-conditional properties of branches in coalescent gene trees

**DOI:** 10.1101/189209

**Authors:** Alexander Platt

## Abstract

Coalescent gene trees have proven to be a powerful framework for formulating and solving problems in population genetics both in theory and practice. Using them, geneticists have been able to generate expectations for many attributes of a random sample of genotypes from a population given a model of the history of the population. This paper derives three new properties of coalescent gene trees that will help characterize the present-day impacts of historical events. Considering a single branch sampled at a given time *t*_*s*_ in the past, it presents distributions describing 1) the length of time a branch sampled *t*_*s*_ generations in the past had existed at the time of sampling, 2) the length of time that branch continues from time *t*_*s*_ towards the present, and3) the the probability that the branch is ancestral to *x* individuals in a modern sample.

## Introduction

The introduction of coalescent theory to the field of population genetics brought with it two key insights: that modeling the genealogy of a sample backwards in time to the point of common ancestry would eliminate the need to keep track of an entire population’s worth of lineages and that properties of the shape of the resulting gene tree would be robust to a wide range of generative evolutionary models1. Where most previous results focus on properties of distributions of times of coalescent events across the entire tree, the properties considered here are those of individual lineages as a function of their positions within a tree. This is accomplished by augmenting the traditional backwards-looking approach to coalescent analysis with forward-looking death models to describe four new properties of coalescent tree branches conditional on their age within the tree. These properties are illustrated in Figure 1, and all are concerned with properties of an arbitrary branch *ℬ* identified in the genealogy of the sample at time *t*_*s*_ before the sample was taken. The first result is the probability density function of *t*_*a*_, the time of the oldest end of branch *ℬ*. The second result is a derivation of the probability density function of *t*_*d*_, the time of the most recent end of branch *ℬ*. The third result is a derivation of the probability mass function of *x*, the number of individuals inthe sample that are descendants of branch *ℬ*, with particular attention paid to the probability that *x* = 1, a special case where branch *ℬ* is considered to be external on the tree. Together, these describe properties of the coalescent tree that are critical to our ability to answer questions about how different parts of the tree relate to each other, how different individuals and groups of individuals in a sample relate to each other, and what kinds of signals historic events may have left in contemporary or archaic samples.

**Figure 1.**
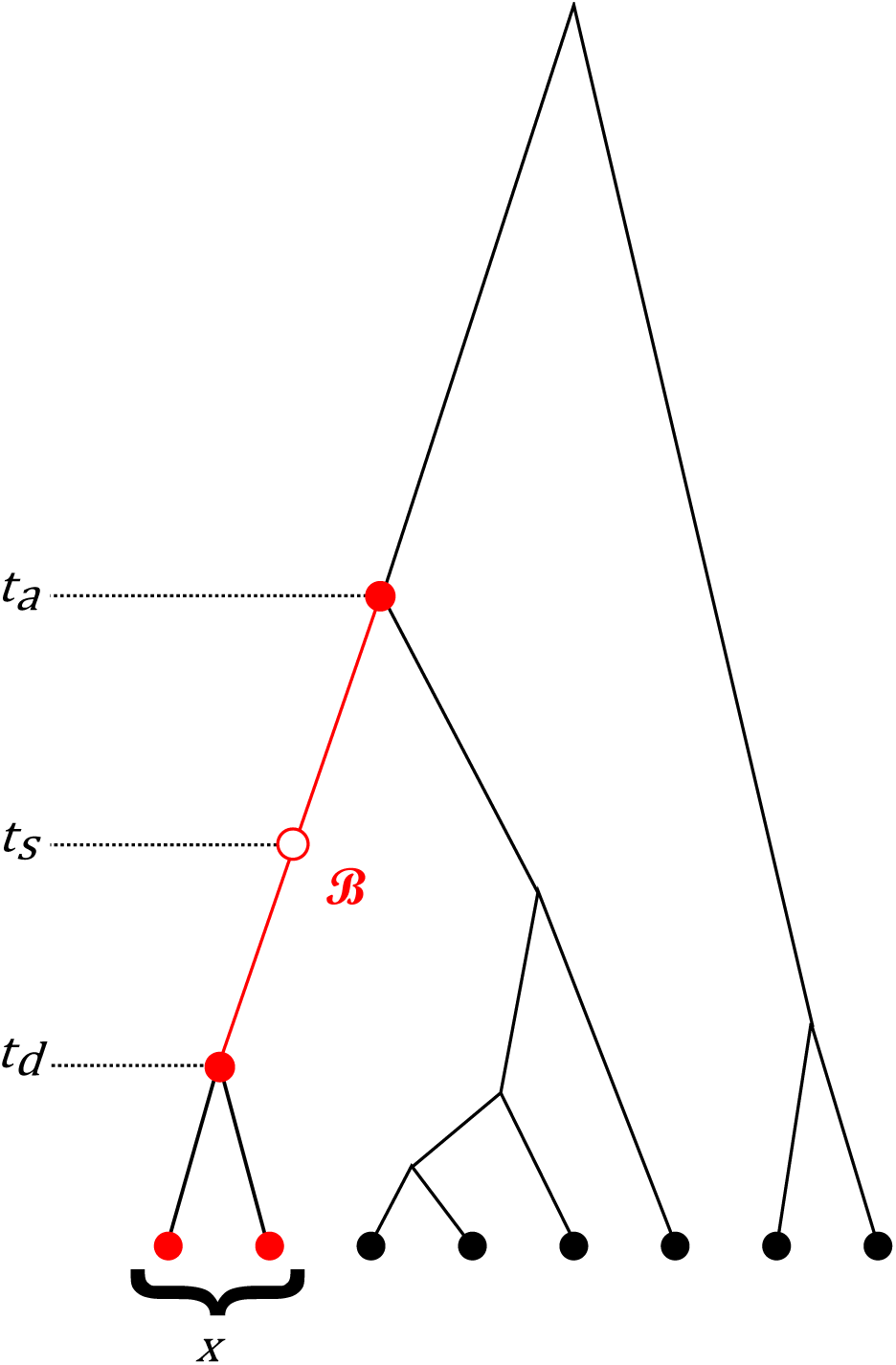
Diagram of important parameters. Branch *ℬ* is identified at time *t*_*s*_. It has an ancestral node at time *t*_*a*_, a descendant node at time *t*_*d*_ and leaves *x* descendants in the current sample.

## Results

### Length of *ℬ* from *t*_*s*_ to *t*_*a*_

Caliebe *et al.*^2^ designate the length of a randomly chosen external branch as *Z*_*n*_0__ in time measured in coalescent units (equal to 2*N* generations for a diploid population of constant size). They do not present a density function for *Z*_*n*_0__ directly but rather the limiting case of the product of the length of a random branch and *n*_0_, the sample size:

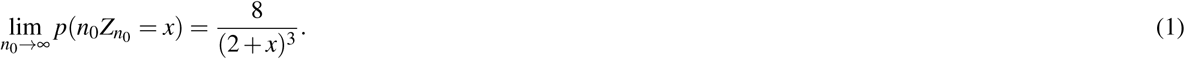

For sufficiently large values of *n*_0_, a change of variable transformation letting 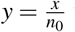 yields

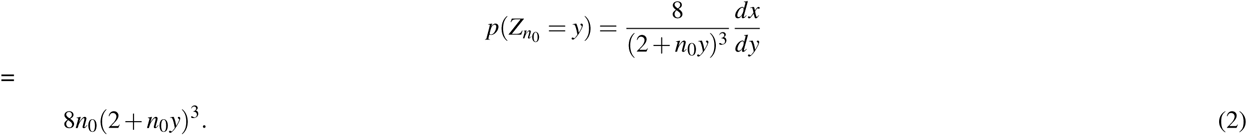

The general distribution of time until the next coalescent event involving a random branch after time *t*_*s*_ is then 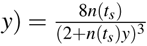 where *n*(*t*_*s*_) is the number of lineages extant at time *t*_*s*_. The full probability distribution for the time-conditional number of lineages is an infinite sum of terms of alternating sign^3^ and is cumbersome to work with. For even fairly small values of *n*, however, it behaves nearly deterministically^4^- and for which numerous approximations exist^5–8^. Most straightforwardly, Slatkin & Rannala6 propose

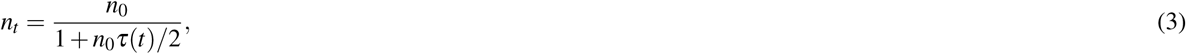

where 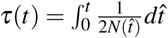 is the coalescent intensity through time *t*^9^ and *N*(*t*) is the historical population size *t* generations in the past.

This approximation produces the result

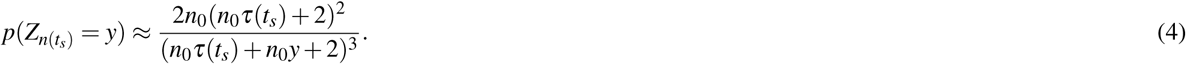

Equations 1, 2, and 4 all yield lengths of time in coalescent units. To convert to generations requires another transformation of variables where *y* = *τ*(*t*_*a*_). This gives

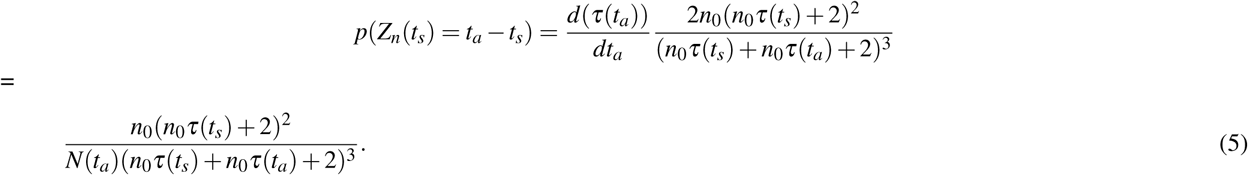

For a population of constant diploid size *N*_*const.*_, this becomes

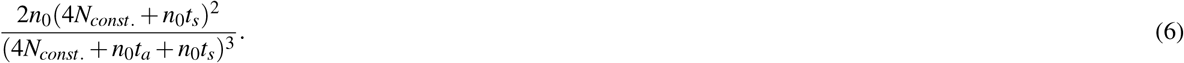

### Length of *ℬ* from *t*_*s*_ to *t*_*d*_

The distribution of the time *ϕ* to the more recent end of a branch identified at time *t*_*s*_ is derived using a hazard model parameterized forwards in time.

The first quantity necessary to derive is the instantaneous rate of coalescence in a population as a function of *ϕ*. Substituting *t* = (*t*_*s*_ − *ϕ*) in equation 3 gives an approximation of the number of lineages extant as a function *ϕ*, and the first derivative of this equation, *nt*(*ϕ*) gives the instantaneous rate of change of the number of lineages through time. As the number of lineages changes by exactly one for every coalescent event, this is equivalent to the instantaneous rate of coalescence:

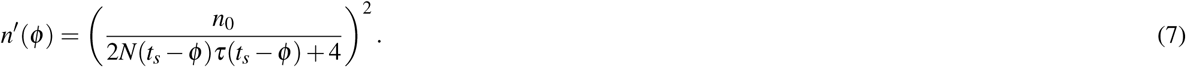

This is the rate of coalescence in the entire population. The only coalescent events of interest, however, are those that would fall along *ℬ*. The probability that any given coalescent event involves *ℬ* is 1*/n*(*ϕ*) as *ℬ* always one of *n*(*ϕ*) lineages. This gives the instantaneous rate of coalescent events on *ℬ* as

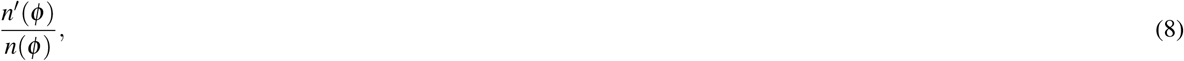

which is referred to as the hazard function *λ* (*ϕ*)^10^. When considering *k* lineages instead of just one, the rate is *k* times faster and

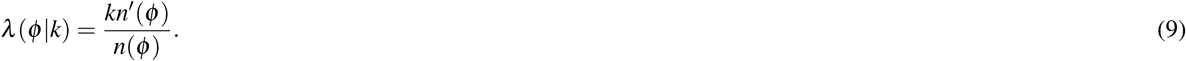

The probability that an event that occurs with an instantaneous rate *λ* (*x*) has not happened over the interval (0, *ϕ*) is 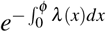 a quantity often referred to as the reliability function or *R*(*ϕ*). The probability that there have been no events on the interval (0*, ϕ*) followed by an event at *ϕ* is simply the product *f* (*ϕ*) = *R*(*ϕ*)*λ* (*ϕ*). Substituting the previous expression for the hazard function gives

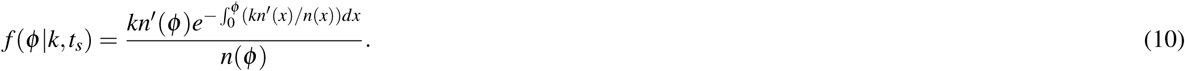

This is the distribution of time until the first coalescent event more recent than *t*_*s*_ within a specified *k* lineages and is true for *ϕ < t*_*s*_. The full distribution gets truncated such that *f* (*ϕ k, t*_*s*_) = 0 when *ϕ > t*_*s*_ and has a point mass at *ϕ* = *t*_*s*_ equal to the *k* = 1 probability from equation (11).

### Distribution of descendants of *ℬ*

A branch is described as external if it the proximal (youngest) end of the branch is a sampled individual, not a coalescent event. The probability that a single branch identified at time *t*_*s*_ is an external branch can be treated as a special case of a more general problem: what is the probability that a set of *k* branches extant at time *t*_*s*_ are all external? Using equation 10 this can be further generalized to the full probability mass function of the number of individuals in the sample descended from *ℬ*.

Consider first the case of *k* = 2 branches identified at time *t*_*s*_, when there are *n*(*t*_*s*_) lineages. The probability that the next coalescent event (forward in time) is *not* among the two selected lineages is

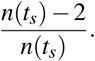

The probability that neither of the first two coalescent events involve the selected lineages is

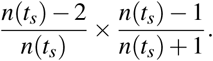

And the probability that none of the first three coalescent events involve the selected lineages is then:

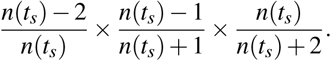

And the probability that none of the first four coalescent events involve the selected lineages is:

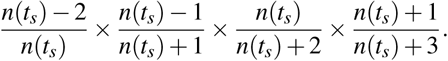

There are in total *n*(*t*_*s*_) *n*_0_ 1 coalescent events between times 0 and *t*_*s*_. The probability that *none* of those events have involved the selected lineages is

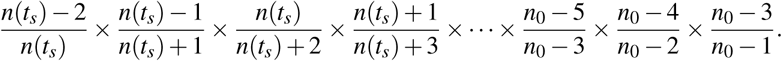

Many of these terms cancel.

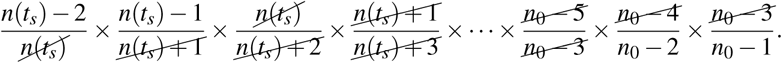

The only remaining terms are the first *k* numerators and the last *k* denominators, leaving the compact probability

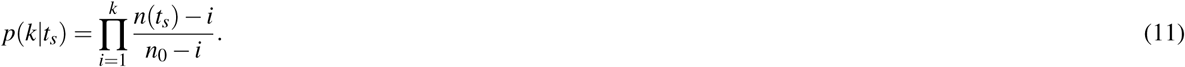

A particular insight from this derivation is the clear observation that *p*(*k|t*_*s*_) depends only on the value of *n*(*t*_*s*_) and not the entire function *n*(*t*). Once we know the number of lineages extant at time *t*_*s*_ it doesn’t matter how or when they got that way. Similarly, the demographic history of the population, *N*(*t*) matters only to the extent that it influences *n*(*t*_*s*_) and is otherwise immaterial.

The distribution of the number of sampled descendants of a branch identified at time *t*_*s*_, *g*(*x* = *X t*_*s*_), is derived by integrating over the possible timing of the intervening coalescent events.

The probability that a branch identified at time *t*_*s*_ leaves exactly one sampled descendant is by definition equivalent to the probability that it is an external branch. The probability that it leaves exactly two sampled descendants is the probability that the lineage splits once at some time *t*_*d*1_, and the two resulting lineages are both external branches. From equations 10 and 11 we get

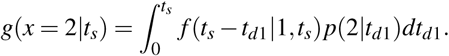

The probability that the branch leaves exactly three sampled descendants is the probability that there have been two coalescent events on lineages descended from *ℬ* at times *t*_*d*1_ and *t*_*d*2_, after which all three resulting lineages are external:

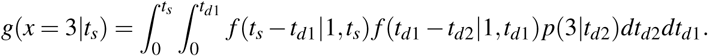

Continuing in this fashion, each successive extra descendant requires integrating over an additional coalescence time and incrementing the number of terminal lineages determined to be external. As a generic function, this gives

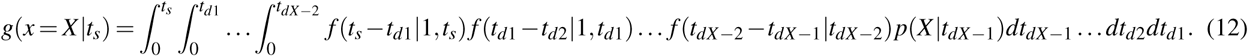

This probability is directly applicable as the number of copies of a variant allele created by a mutation at time *t*_*s*_ (conditional on the mutation being present in a sample). In general, this expression will be of most use for problems involving branches with relatively few descendants, where the multiplicity of integrals (there are *X -* 1 of them) won’t be burdensome.

## Discussion

By using standard coalescence theory to describe the overall shape of a gene tree and forward-in-time death process analysis it is possible to get simple closed form expressions related to arbitrary branches within the tree as functions of the time at which those branches existed.

## Acknowledgements

With thanks to Jody Hey for helpful comments and discussion, and funding from NIH Grant RO1GM078204 to Dr. Hey.

